# Uncovering putative neural mechanisms of neurotherapeutic impacts on EEG using the Human Neocortical Neurosolver

**DOI:** 10.64898/2026.04.11.717895

**Authors:** Nicholas Tolley, David W. Zhou, Austin E. Soplata, Dylan S. Daniels, Katharina Duecker, Carolina Fernandez Pujol, Joyce Gao, Stephanie R. Jones

## Abstract

A key barrier to developing effective drugs for disorders of the central nervous system (CNS) is understanding their impact on neural circuits. This protocol demonstrates how physics-based neural simulations can be used to interpret electrophysiological biomarkers of neurotherapeutics, providing a mechanistically grounded approach to the development of neurotherapeutics.

**LONG ABSTRACT:** Electroencephalography (EEG) and electrophysiology methods provide millisecond resolution biomarkers for central nervous system disorders and are used to assess treatment-related effects. However, lack of understanding about the neural mechanisms generating such biomarkers impedes the development of diagnostics and therapeutics based on these signals. The Human Neocortical Neurosolver (HNN) is an open-source biophysical modeling software that connects localized EEG biomarkers to their multi-scale neural generators. This protocol demonstrates a hypothesis-driven workflow using HNN to test possible neural mechanisms of neurotherapy-induced EEG biomarkers by optimizing parameters to achieve a fit between simulated and empirical current source waveforms. Corresponding multi-scale cell- and circuit-level activity can then be visualized and quantified, providing validation targets for model predictions in follow up empirical studies. An example is provided which shows how to examine the generating mechanisms of the early event-related potential (ERP) components of an auditory evoked response (P1, N1 and P2) and to assess changes following neural circuit modification due to neurotherapeutic administration. This protocol demonstration enables scientists to design simulation experiments to develop testable predictions on how EEG biomarkers reflect neural circuit mechanisms of example therapeutics. A similar protocol can be applied to study disease mechanisms or other therapies.

## INTRODUCTION

Central nervous system (CNS) therapeutic development suffers from unique challenges, with approval rates lower than those seen in other disease areas, spurring the need for innovative methodological approaches^1^, particularly those that can uncover treatment-related effects on brain dynamics. A well-established approach to study the impact of therapeutics on neural activity is electroencephalography (EEG)^2, 3^. EEG provides a signature of *in vivo* circuit-level brain dynamics and offers promise for translating rodent models to human trials, as the neural circuitry that generates EEG bears homology between mice and humans^4–8^. In pharmaceutical development, EEG studies can potentially serve multiple important functions: providing translational readouts to bridge animal and human trials, evaluating drug safety, guiding lead compound selection, informing dose-response relationships, assessing proof-of-mechanism in early clinical phases, enabling clinical trial stratification and cohort enrichment, and more^9–14^.

A robust EEG biomarker used in CNS drug discovery is the event related potential (ERP, see Supplementary Table 1 for a list of all acronyms used in this manuscript). ERPs reflect time-locked sensory evoked brain activity and have been used to study neurodevelopmental and neuropsychiatric diseases, including depression^15, 16^, schizophrenia^17, 18^, autism-spectrum disorder^19, 20^, and Alzheimer’s disease^21^. They have also been used to study the impact of CNS drugs on brain circuits^22, 23^ where treatment-induced normalization toward healthy responses may indicate therapeutic efficacy^24^. A limitation of ERPs, and other standard EEG biomarkers, (e.g. brain oscillations) is that while their features can be associated with disease states or drug effects (e.g. dose ranges^25, 26^), these associations are correlational in nature, and the causal influence of cell and circuit activity on the biomarker generation is unknown. Understanding the cell and circuit origins of EEG biomarkers can potentially deepen the value of EEG by mechanistically linking signatures to underlying physiology (also see^27, 28^, note that we use the term EEG “biomarker” to refer to measurable EEG changes with therapeutic intervention, consistent with the FDA NIH BEST framework definition^29^, rather than implying that these signals are formally qualified for a specific context of use^30^). Technically sophisticated invasive recordings can provide detailed cell- and circuit-level interpretation of EEG biomarkers, however they are largely restricted to animal models. Instead, biophysical simulations offer a principled and powerful framework for bridging this gap by simulating the physical processes through which neural circuits produce measurable EEG signals (Figure 1).

**Figure 1:**
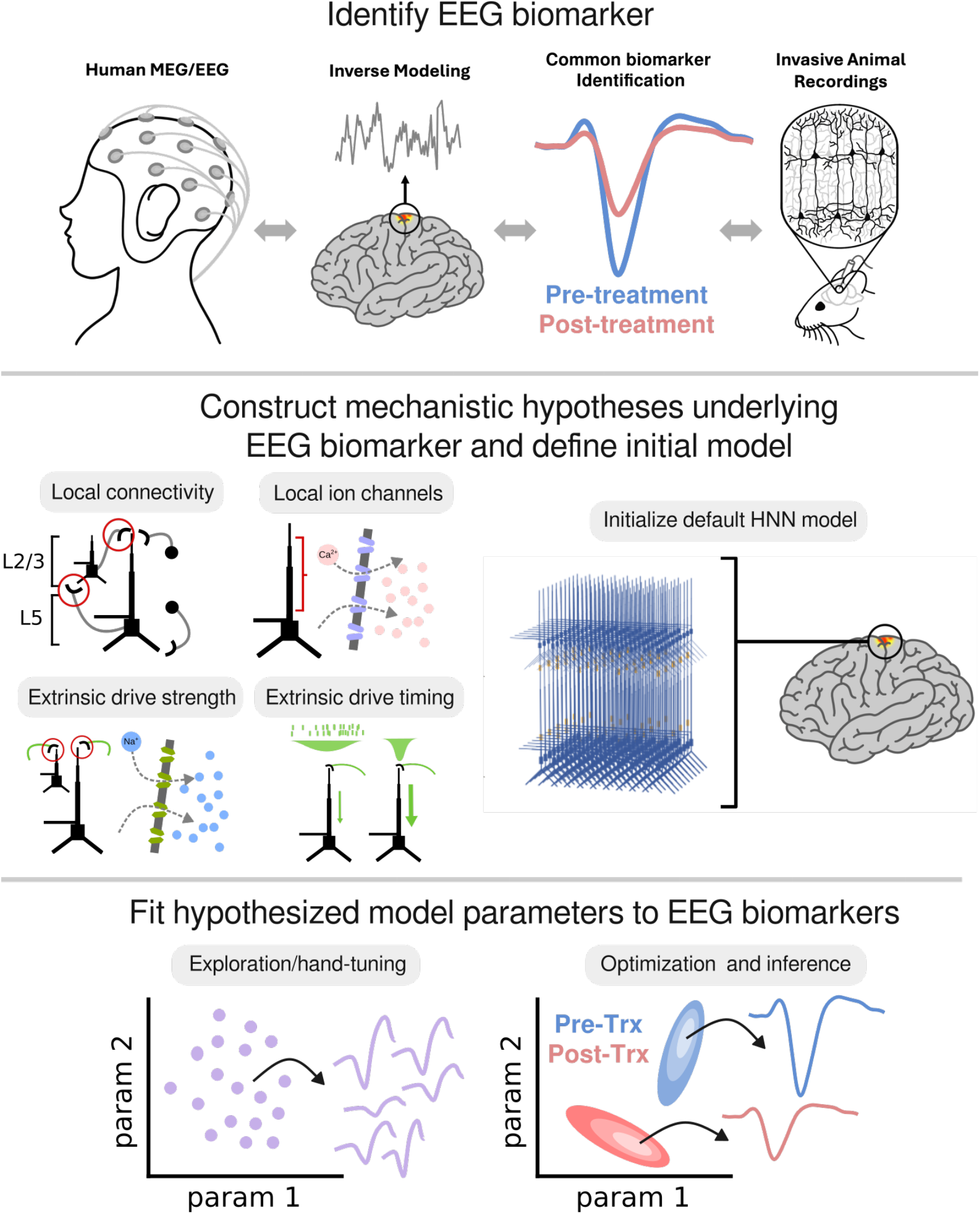
Biophysical modeling to develop and test mechanistic hypotheses underlying pharmacological EEG biomarkers. Top: Modeling EEG biomarkers begins with picking a specific brain signal that is reliably different between patient populations. An example of a hypothetical EEG biomarker is an auditory event related potential (ERP) that is suppressed in post-treatment (red) relative to pre-treatment (blue). Middle: Biophysical modeling allows for testing mechanistic hypotheses that explain how EEG biomarkers emerge and change with drugs. Hypotheses about which drug mechanisms lead to distinct brain activity patterns must be constructed, and corresponding model parameters identified. Bottom: The default HNN model is used as a starting point to test hypotheses by either manually altering the values of the chosen model parameters, or using automated optimization and inference algorithms. Differences in parameter values pre-to post-treatment correspond to model-based predictions.

Several modeling frameworks exist for interpreting the neural generation of EEG, with varied degrees of model complexity^31–34^. This protocol uses the biophysical modeling framework of the Human Neocortical Neurosolver (HNN) to link ERP biomarkers of treatment-related effects to their underlying cell- and circuit-level mechanisms^33^ (Figure 2). HNN’s architecture is based upon the principle that synchronous intracellular current flow of aligned pyramidal neuron dendrites generates the primary electrical current sources (i.e. current dipoles) underlying EEG signals^6, 35–37^. HNN has been designed and distributed with workflows and default parameter sets to simulate the origin of several commonly studied EEG signals, including ERPs. HNN’s functions also include methods for direct statistical comparison to empirical data, and is constructed with a modular design to facilitate the addition of new cell and circuit features as new knowledge suggests they are necessary.

**Figure 2:**
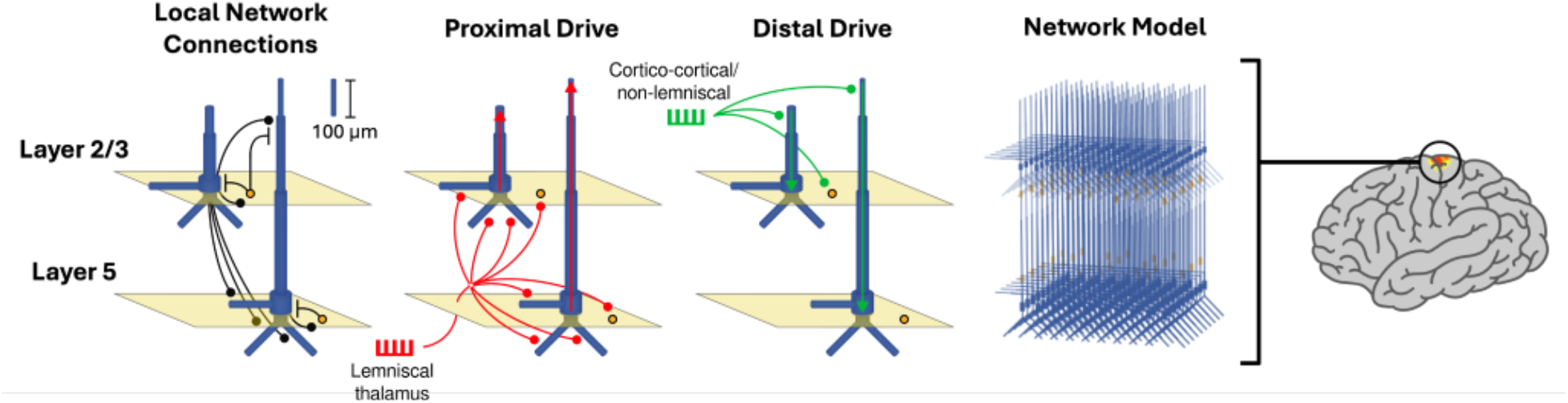
Schematic of the Human Neocortical Neurosolver (HNN) model. Visualization of the major components of the HNN model, namely the local network connections between excitatory and inhibitory neurons, and the exogenous input pathways named “proximal drive” and “distal drive”.

HNN simulations may be applied at key decision points throughout the pharmaceutical R&D workflow, from interpreting pharmacodynamic brain signatures to designing mechanistically informed biomarker strategies^14, 38–40^. Potential use cases include: target validation, comparing drug mechanisms of action, dose finding, targeting therapeutic modulation of a specific circuit mechanism, and generating hypotheses for follow-up experiments. This protocol specifically focuses on HNN simulations of the early P1, N1, and P2 components of a simulated auditory ERP due to extensive literature characterizing the neural mechanisms of these early ERP components, which helps constrain hypothesis-driven modeling ^41^. While we focus on drug-induced ERP biomarkers, the protocol can be adapted to study other neurotherapeutics (e.g. brain stimulation and biofeedback training), or mechanisms of CNS disorders (e.g. depression and schizophrenia).

#### Box 1: Overview of HNN network model

All biophysical neural models are a reduced representation of the full complexity of the brain. The level of biophysical detail chosen for HNN was designed to account for the biophysical generation of the primary current sources underlying localized EEG (or magnetoencephalography (MEG)) signals, known to emerge from the intracellular current flow in long and spatially aligned cortical pyramidal neuron dendrites (Hämäläinen et al., 1993; Ikeda et al., 2005; Murakami & Okada, 2006; Okada et al., 1997), together with other cell and circuit elements that critically influence pyramidal neuron dynamics (e.g. inhibitory neurons, and synaptic connections from exogenous networks). As such, the HNN model (Figure 2) is composed of excitatory pyramidal neurons, and inhibitory interneurons, distributed in a 2 layer network. The default HNN template model contains 100 pyramidal neurons synaptically coupled to 33 inhibitory cells in both layer 2/3 (L2/3) and layer 5 (L5) (266 neurons total) representing a canonical neocortical column structure in a small patch of neocortex. A multiplicative scaling factor is used to estimate the signal from a larger synchronous network. To account for dendritic morphology, pyramidal cells are simulated with a small number of multicompartment sections representing characteristic dendrites^42^, while inhibitory cells are modeled as single compartments given their negligible contribution to recorded currents^33^. All cells contain active ionic currents in the somatic and dendritic compartments that are simulated with Hodgkin-Huxley style dynamics. Pyramidal cells connect to other cells with both excitatory AMPA and NMDA synapses, while basket cells connect with inhibitory GABA_A_ and GABA_B_ synapses. To simulate ERPs, or other EEG biomarkers such as oscillations, the network is activated by excitatory synaptic drive from exogenous brain areas that innervate the network through layer-specific canonical connection pathways. One pathway corresponds to feedforward sensory input from lemniscal thalamus that targets layer 4 and then propagates to the proximal dendrites of the pyramidal neurons and interneurons, and is referred to as “**proximal drive**”. The other pathway corresponds to feedback information from other cortical regions, as well as non-lemniscal thalamus, that directly targets distal pyramidal neuron dendrites and interneurons in supragranular layers, and is referred to as **“distal drive”**. The exogenous brain networks providing these drives are not explicitly simulated, but rather represented by user-defined trains of action potentials that activate the network through the proximal and distal projection pathways and induce excitatory postsynaptic currents and spiking interactions that generate intracellular dendritic current flow up and down the aligned pyramidal cell dendrites. The net simulated current flow across the population of pyramidal neurons is expressed in units of nano-ampere-meters (nAm) and can be one-to-one compared to orientation-constrained source localized E/MEG signals, which are also computed in nAm units. The additional multi-scale detail included in HNN provides targets for model validation and/or updating through invasive recordings in animals^7^, or with other imaging modalities^43^.

The parameters and ERP workflow in the default canonical HNN model were informed by available data and initially constrained with information from the primary somatosensory cortex (see^44–46^). An ERP is simulated by activating the local network with a predefined sequence of exogenous thalamocortical and cortico-cortical drives evoked by sensory stimulation. This drive sequence consists of an initial feedforward proximal drive (e.g. see leftmost red histogram near the time of the P1 in Figure 4E below), followed by a feedback distal drive (see green histogram near the time of the N1 in Figure 4E), followed by a re-emergent feedforward proximal drive (see red histogram near the time of the P2 in Figure 4E rightmost red histogram). The parameters defining these driving inputs were constrained to produce a close fit to source localized primary somatosensory ERPs in healthy human adults emerging from brief finger taps. However, due to the canonical structure of cortical circuity, the default HNN framework has proven useful to studying signals from other brain areas including auditory^47–49^, visual^50^, and in frontal cortex^51^), and model-derived predictions have been validated with empirical recordings in follow up studies^7, 43, 46^. Please see^33^ for further details of default network construction.

### Overview of iterative process for using HNN as a hypothesis development and testing tool

Large multi-scale neural models such as HNN contain thousands of parameters, whose values are informed by empirical data to the extent possible. However, many initial assumptions are made and the use, validation, and updating of such models is an iterative process of modeling and experimentation. The process begins with data-constrained model development (i.e. define structure and initialize parameters) using currently available data and/or information from literature. Once an initial model is constructed, a targeted subset of parameters can be fit to empirical data, and uncertainty estimates can be generated. Due to the multi-scale nature of the model, simulations constrained to one (or more) types of data generate predictions about the full system dynamics, providing multiple targets for empirical validation. For example, in the ERP protocol described here, an initial pre-tuned HNN model is provided and a subset of parameters (referred to as “**parameters of interest**”) are adjusted to create a close match between simulated and empirically recorded primary current dipoles. Once fit to the empirical current dipole, predictions on all other cell and circuit dynamics are simultaneously produced (e.g. cell spiking activity, layer specific local field potentials (LFP) and current source density (CSD)). Each of these generative predictions provides targets for model validation in empirical studies. If the predictions are falsified by experiments, then the model is updated by constraining it to the new data (Figure 3).

**Figure 3:**
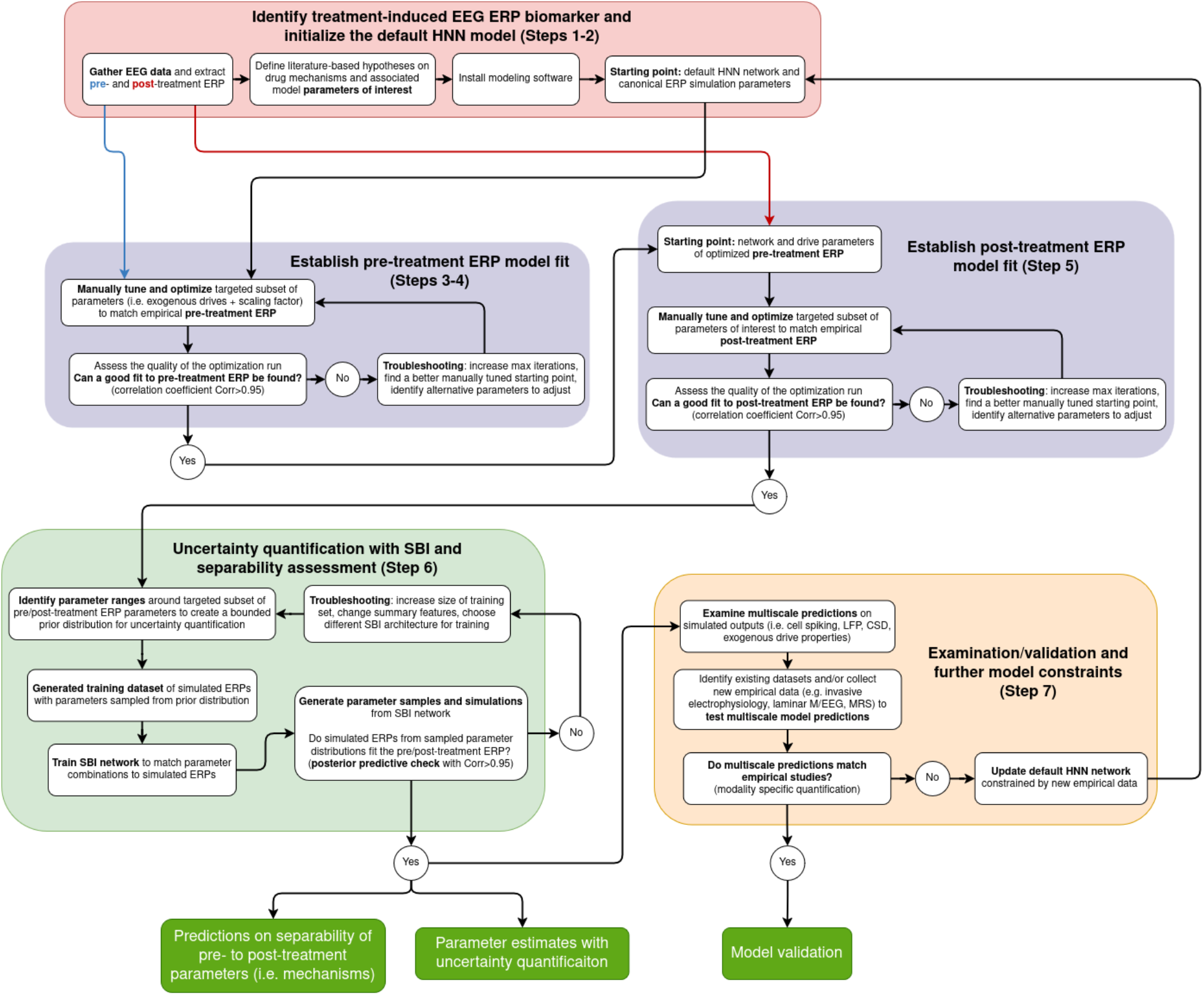
Iterative workflow for developing and testing ERP biomarker predictions with HNN. The flow chart maps onto the steps of the protocol, beginning with “Identify treatment-induced EEG ERP biomarker and initialize default HNN model” (red box, **Steps 1-2**). Optimization and hand-tuning is then performed to fit model parameters to the pre-treatment and post-treatment ERP signals (purple box, **Steps 3-5**). Uncertainty quantification of these fit model parameters is then performed with SBI (green box, **Step 6**), and lastly, predictions from the model can be examined and compared with experimental data (e.g. invasive electrophysiology) to validate model-derived predictions or provide further constraints to update the multi-scale model (orange box, **Step 7**).

### Iterative workflow for developing and testing ERP biomarker predictions with HNN

(Figure 3). In the current protocol, the initial pre-tuned default HNN model described above (distributed with the software) is used as the starting point for examining pre-to post-treatment ERP biomarkers (Figure 3 **Steps 1-2**). With a pre-to post-treatment ERP biomarker identified (e.g. post-treatment changes in ERP peaks and latencies), users must define a limited subset of target **parameters of interest** hypothesized to be impacted by the drug based on prior studies and literature. Experimental studies on drug-induced changes at the level of ion channels and synapses are particularly relevant for parameter selection (e.g.^52, 53^). In this protocol, we give example post-drug parameters to test, including parameters for the exogenous drive timing and strength, local inhibitory connectivity, and dendritic ion channel conductances.

Beginning with default ERP simulation parameters, users first fit a limited set of **parameters of interest** to match simulated ERPs with the pre-treatment empirical ERP. In this protocol, we provide the **parameters of interest** to start with, namely the scaling factor and parameters of the exogenous drive. The fitting process begins by manually adjusting (i.e. “hand tuning”) the scaling factor parameter to better fit the overall magnitude of the canonical ERP simulation to the empirical ERP (Figure 3 **Steps 3-4**). Next, the timing and the strength of the exogenous sequence of proximal and distal driving inputs are tuned to match the timing and magnitude of the peaks in the empirical pre-treatment ERP. Once an initial close representation to the data is found with hand tuning, automated parameter estimation is used to improve the fit. Several automated parameter fitting procedures are distributed with HNN, including covariance matrix evolutionary strategy (CMA-ES), Bayesian optimization, and constrained optimization by linear approximation (COBYLA). These automated approaches search the parameter space to find a single set of values that produce the closest match to the target EEG data. Once a pre-treatment simulation is achieved, the **parameters of interest** hypothesized to account for the post-treatment ERP are also fit using the parameterized pre-treatment simulation as a starting point (Figure 3 **Step 5**).

With both the pre- and post-treatment model fits established, SBI is used to perform uncertainty quantification of the fitted parameters (Figure 3 **Step 6**). In large-scale biophysical models, the parameters fit through optimization are not guaranteed to be uniquely good solutions. SBI is used to estimate distributions of model parameters that all account for pre- and post-treatment ERPs. Separability of the parameter distributions correspond to predictions on mechanisms of action through which the neurotherapeutic distinctly impacts cortical circuitry, and the degree of separability can be quantified using a distribution overlap index (OVL^54, 55^). From there, a detailed examination of the underlying cell and circuit activity can be visualized and quantified, providing multi-scale targets to validate model derived predictions (Figure 3 **Step 7**), or further constrain and update the model when predictions are negated. The scope of this protocol is restricted to the generation of multiscale predictions. Specific techniques to quantify and validate different measures (e.g. cell spiking) are not explicitly provided. We note that while **Steps 3-6** describes fitting the model to ERP data, the fitting procedure described can be readily adapted to other data types, as new empirical data becomes available (e.g. cell spiking, LFP, and CSD).

#### When to use this protocol

This protocol is appropriate for interpreting orientation-constrained preprocessed and source-localized E/MEG data collected during an evoked response paradigm (ERPs). Many frameworks exist to perform data cleaning (i.e. preprocessing) and source localization (e.g. MNE-Python^56^). The necessary data format to import to HNN is source-localized data, with nano-ampere-meter (nAm) units, saved in a time-vector aligned to the stimulation onset time. In the case of fast sensory evoked responses, source- and sensor-level time courses are highly analogous and knowledge gained from studies of source-localized data can be applied to interpret sensor-level data (e.g. see^57, 58^).

### PROTOCOL

The dataset used in this research was extracted from a previously published study^48^. As the data is from a published source, separate ethical approval was not required for this manuscript.

1. **Identify a treatment-induced EEG ERP biomarker** (i.e., pre-to post-treatment change) for biophysical modeling and form hypotheses about model parameters representing treatment-related effects (i.e., **parameters of interest**). *NOTE: EEG biomarker characterization is expected to be completed prior to biophysical modeling. This protocol does not fully describe the preprocessing necessary for EEG-based biomarker characterization. Several prior works describe the preprocessing and analysis of EEG signals in full detail; readers are particularly invited to check out*^*56*, *59*^ *for a more complete background*.
  1.1. Collect or identify dataset with experimentally recorded EEG signals from subjects of interest (e.g. pre-treatment and post-treatment in the context of neurotherapeutics). ERP experiments require EEG measurements recorded during presentation of a sensory stimulus; time stamps of sensory stimulus must be recorded simultaneously with EEG data for segmentation into trials.
  1.2. Identify a set of candidate ERP biomarker features that are hypothesized to distinguish treatment-related effects (e.g. ERP peak timings and magnitudes). *NOTE: In this example protocol we use peak magnitudes as a biomarker of interest*.
  1.3. Preprocess EEG data and extract biomarker features of interest. *NOTE: Several software packages support the preprocessing and ERP analysis of EEG data including MNE-Python*^*56*^, *EEGLab*^*60*^, *and FieldTrip*^*61*^. *While source-localization is assumed in this protocol and recommended for modeling ERP signals, this is not a required step (see Box 1). An example of source localization can be found at https://jonescompneurolab.github.io/hnn-core/stable/auto_examples/workflows/plot_simulate_somato.html*.
    1.3.1. Perform source localization using sensor-level signals from all channels, or choose EEG sensors to be analyzed *NOTE: One-to-one unit correspondence shown below will not hold for sensor signals*.
    1.3.2. Segment recorded EEG data into trials using timestamps of sensory stimulus.
    1.3.3. Compute trial-averaged ERP waveform for both pre-treatment and post-treatment conditions.
    1.3.4. Extract candidate ERP biomarkers from trial-averaged waveforms (e.g. compute N1 peak magnitudes).
  1.4. Conduct statistical tests to determine which ERP features are significantly different across conditions (e.g. pre-vs-post-treatment). *NOTE: A code example of statistical tests applied to ERP features can be found at https://mne.tools/stable/auto_tutorials/stats-sensor-space/20_erp_stats.html*
  1.5. Output: Specific statistically significant distinguishing EEG biomarker features (e.g. difference is N1 magnitudes).
  1.6. **Define literature-based hypotheses on drug mechanisms and associated model parameters of interest**. Consult literature and experimental data to identify biophysical properties that are presumed to be altered by the neurotherapeutic under investigation that may account for feature differences.
  1.7. Identify which parameters of the biophysical neural model (i.e. HNN) are either directly represented, or indirectly related to the biological properties identified in **Step 1.6**. These are known as **parameters of interest**.
  1.8. Output: An identified set of model parameters of interest that correspond to biophysical properties that are hypothesized to generate identified EEG feature differences. The default HNN (initialized in **Step 2**) provides an initial value for all parameters.
2. **Initialize the default HNN model:** Install modeling software and set up the project folder. *NOTE: The software versions used in this study are specified in the Table of Materials, along with minimum system requirements. Multiple installation options are available (i.e. pip, conda, and source installation), for Linux, MacOS, and Windows*.
  2.1. Download and install a functioning version of Anaconda Python.
  2.2. Install the HNN-core software using the operating system specific instructions (https://jonescompneurolab.github.io/textbook/content/01_getting_started/installation.html). *NOTE: To efficiently install the software dependencies used in this study, the associated code repository (*https://github.com/ntolley/hnn_jove*) uses pixi (https://pixi.prefix.dev/latest/). Instructions for installation of pixi and a local installation of the code repository are located on the repository’s README.md file*.
  2.3. Verify installed version of HNN-core is 0.6.0 or greater by typing the following into the terminal: **pip show hnn_core**
  2.4. Launch the HNN-GUI by typing **hnn-gui** in the terminal and hitting enter.
  2.5. Create a new project folder in the computer’s file system where all data files generated in this protocol will be stored.
3. **Establish pre-treatment model fit with manual tuning**. Starting with canonical HNN ERP simulation and its default parameters, manually tune scaling factor and exogenous drive parameters to fit pre-treatment ERP (e.g. pre-treatment ERP). *NOTE: The HNN graphical user interface (GUI) automatically loads model parameters fit to a somatosensory event related potential*^*45*^, *which through numerous studies has been shown to be a good “canonical ERP” starting point. This tutorial will focus on modifying the scaling factor and exogenous input parameters from this starting point*.
  3.1. Load the pre-treatment empirical ERP waveform from **Step 1** into the HNN GUI (Figure 4, orange waveform).
    3.1.1. Click the **Load data** button on the menu bar located on the lower left portion of the GUI window (Figure 4D). *NOTE: The nomenclature on ERP peak naming is widely varied across literature, as such, the P1/N1/P2 labels in Figure 4F which are for illustrative purposes only and may not correspond to the same naming conventions used in other studies*.
    3.1.2. In the file browser window, select a .**csv** or.**txt** file containing the ERP waveform to be modeled (i.e. the target waveform). Files must be formatted with 2 columns, where the first column contains the time in ms, and the second column contains the source-localized empirical dipole waveform in units of nAm. *NOTE: The empirical data file is named pre-treatment.txt in the associated code repository*.
    3.1.3. Inspect the waveform that is automatically plotted in the figure panel (Figure 4F).
  3.2. Run the default simulation of a canonical ERP.
    3.2.1. Set the parameter values of **tstop, dt, Trials, Backend**, and **Cores** in the **Simulation Parameters** panel (Figure 4B) to the desired values. The **tstop** parameter controls the simulation length, **dt** controls the integration time-step, and the **Trials** parameter controls the number of repeat simulations run with the same model parameter values. **Backend** is a choice between serial (Joblib) and parallel (MPI) simulations, run over a chosen number of computer **Cores**. *NOTE: Variability across trials comes from the Std. dev time of the exogenous evoked drives described in* ***Step 3.5*** *below*.
    3.2.2. Click the **Run** button (Figure 4D) to start the default simulation of a canonical ERP.
  3.3. Create a plot that compares simulated ERP from **Step 3.2** to the loaded empirical pre-treatment ERP. *NOTE: The HNN-GUI provides the option to calculate two different goodness of fit measures: correlation coefficient (Corr) and root mean squared error (RMSE). The goal of manual hand tuning and optimization (****Step 4***).
    3.3.1. Click the **Visualization** tab on the top left of the GUI window (Figure 4A).
    3.3.2. Click the dropdown menu labeled **Data to compare** (not shown) to select the loaded target waveform from **Step 3.1**.
    3.3.3. Click **Clear axis** to reset the plot.
    3.3.4. Click **Add plot** to generate a new plot with the simulated initial ERP waveform (blue) and target waveform (orange) overlaid, along with text indicating an automatically calculated correlation coefficient and RMSE between these two waveforms (Figure 4F).
  3.4. Modify with manual hand tuning the scaling factor to approximately match the magnitudes of the simulated and empirical dipole waveforms. Set the default **Dipole scaling** parameter (Figure 4C) on the **Simulation tab** (Figure 4A). *NOTE: The scaling factor corresponds to a prediction on the estimated number of neurons underlying the generation of the EEG signal. The default of 3000 indicates that 200 pyramidal neurons (size of HNN model) x 3000 = 600,000 neurons are necessary to generate an evoked response with the magnitude in nAm indicated on the y axis of Figure 4F*.
  3.5. Modify with manual hand tuning the mean and standard deviation of exogenous drives to get a closer fit to the timing of the empirically recorded pre-treatment ERP peaks (i.e. P1/N1/P2) (Figure 5 and Figure 7A to Figure 7B). *NOTE: The default local connectivity and cell parameters distributed with HNN were tuned to reproduce healthy single-cell and network-level activity patterns. While the local network parameters can also be adjusted, in practice it is recommended that users leave the pretuned local HNN neocortical template model parameters fixed, and first test if a reliable fit to the data can be found by only adjusting the exogenous drives*.
    3.5.1. Identify which simulated ERP peaks are misaligned in time with the empirical ERP waveform (Figure 4). *NOTE: This example assumes there are three early peaks in the* empirical *ERP, as in the default canonical ERP simulation. To add peaks, additional external drives can be simulated*.
    3.5.2. Click the **External drives** tab on the top left of the GUI window (Figure 4A and Figure 5A). *NOTE: The parameters for three predefined exogenous drives will be visible, representing the feedforward proximal (evprox1), feedback distal (evdist1), and re-emergent feedforward proximal drive (evprox2) generating the default canonical ERP simulations (see Box 1). Histograms depicting the spike times and counts of these default drives are shown in Figure 4E*.
    3.5.3. Click on the drop down of the exogenous drive whose **Mean time** is closest to the misaligned peak.
    3.5.4. Modify with manual hand tuning the values in the text boxes for **Mean time** and **Std dev time** to better match the timing/width of peaks in the target waveform (Figure 5B-D). The **Mean time** can be adjusted to shift the peak time, the **Std dev time** can be adjusted to change the width of the peak. *NOTE: The Mean time and Std dev time control the mean and variance of the exogenous spikes that activate the local network in proximal or distal project patterns (see histograms in Figure 4E). These parameters do not fully control the timing/width of ERP peaks. The exact timing and width of ERP peaks depends on both exogenous drives and intrinsic activity of the local network. Figure 5 shows changing the* ***Mean time*** *to align with the ERP peaks at 60 (evprox1), 100 (evdist1), and 150 (evprox2) ms*.
  3.6. Modify with manual hand tuning the synaptic weights (i.e. post-synaptic conductance) of exogenous drives (Figure 6) to get a closer fit to the magnitude of the empirically recorded ERP peaks (i.e. P1/N1/P2) (Figure 7B to Figure 7C)
    3.6.1. Identify which simulated ERP peaks are misaligned in magnitude with the empirical ERP waveform.
    3.6.2. Click the **External drives** tab on the top left of the GUI window (Figure 4A).
    3.6.3. Click on the drop down of the exogenous drive whose **Mean time** is closest to the misaligned peak.
    3.6.4. Modify the values in the text boxes under **AMPA weights** and **NMDA weights** to adjust the synaptic conductances. Larger proximal drive strength to L5 and L2/3 pyramidal neurons generally produces more positive peaks, and distal drive strength generally produces more negative peaks. *NOTE: Similar to the exogenous drive timing, the magnitude of ERP peaks is not fully determined by the strength of exogenous drives. Spiking dynamics evoked by the drives can lead to changes in peak magnitudes that are difficult to predict a priori. It is recommended to test one order of magnitude changes in strength to start (e.g. AMPA L5_pyramidal 0.014 to 0.14) and examine the response, then adjusting up or down from there. Figure 6 shows changing values to be 10x smaller than the default canonical simulation*.
  3.7. After all desired changes in **Steps 3.4-3.6** have been made, click the **Simulation** tab (Figure 4A) and type a new name for the simulation in the **Name** text box (Figure 4B)
  3.8. Click the **Run** button to simulate the modified parameter set. Once the simulation is completed, a new plot will be generated in the Figure panel (Figure 4F, see also Figure 7). Previous plots can be accessed by clicking the corresponding figure tabs (i.e. the tabs labeled “Figure 1” and “Figure 2” above the panel Figure 4E).
  3.9. Continue iterative manually tuned modification to attempt to further increase the correlation coefficient. Repeat **Step 3.3** to replot the simulation with the target waveform, and recalculate the correlation coefficient.
  3.10. Click the **Save Network** button to save the best fit parameter set in a .**json** file, and the **Save simulation** button to save a .**txt** file containing the simulated dipole wavework (Figure 4D). Move both files to the project folder created in **Step 2.5**. The file names will match the name of the simulation in the dropdown menu. *NOTE: All files are saved to the default download directory of the web browser used to run the GUI. It is recommended to either 1) manually move the files to the project folder created in* ***Step 2.5***, *or 2) temporarily change the default download directory of the web browser running the GUI*.
4. **Establish pre-treatment model fit with parameter optimization**. Apply automated parameter optimization to exogenous drive parameters to better fit the pre-treatment ERP (ie. pre-treatment ERP). *NOTE: This example shows how to optimize targeted parameters to estimate single values that produce a close fit to the waveform using CMA-ES (not to be confused with SBI; both are approaches to fit model parameters but the primary output of SBI is a distribution). An example of how to estimate distribution of parameters that can account for waveforms is given in Figure 10. For pre-treatment ERPs, it is recommended to start by optimizing the exogenous drive parameters under the assumption that the cell and local network connection parameters in the default HNN neocortical model are fixed. The multiscale prediction provided by HNN described in* ***Step 7****provides targets for validation of this assumption and, as new information to constrain model predictions becomes available, the flexible HNN framework allows for estimation of any set of parameters*.
  4.1. Click the **Optimization** tab on the top left corner of the GUI (Figure 8A).
  4.2. Configure the settings of the optimization run (i.e. number of iterations, solver, and objective function). *NOTE: The default optimization settings (****Objective function****=“dipole_corr”;* ***Solve****r=“cma”) are appropriate for ERP waveforms. This default Objective function maximizes the correlation coefficient between the simulated and empirical waveform. The* ***Max iterations*** *may need to be increased if many parameters are optimized. The correlation coefficient is a scale-free measure, therefore when using “dipole_corr”, the scaling factor will need to be adjusted after optimization (****Step 4.8.1****). Alternatively, users may choose “dipole_rmse” for the objective function, which minimizes the RMSE between the simulated and empirical waveform, and in this case the scaling factor remains fixed*.
    4.2.1. Click on the **Max iterations** textbox and enter 100
  4.3. Click on the dropdown menu of an exogenous drive whose parameters will be optimized (Figure 8A-B red circle).
  4.4. Select the drive parameters to be optimized by clicking the checkbox under **“Optimized against?**” (Figure 8B).
  4.5. Specify the range of parameter values that can be explored by the optimizer by typing the desired values into the **Min** and **Max** textboxes under **Constraints (in %)**(Figure 8B). *NOTE: The default values of 20% are suitable small changes for a simulation that already has a high correlation coefficient Corr > 0.9 (e.g. in Figure 8B a range of 20% applied to the* ***Mean time*** *of 65.53 ms produces bounds of 52.42-78.64 ms). Optimizing waveforms with a poor initial fit can potentially be achieved by increasing the Min and Max percentages, however, the number of simulations required to find a correlation coefficient may significantly increase*.
  4.6. Run the optimization routine by clicking the **Run Optimization** button (Figure 8A).
  4.7. Save the optimization run’s history by clicking the **Save Optimization History** button (Figure 8A). Move the file to the project folder created in **Step 2.5**.
  4.8. Assess the quality of the optimization run. *NOTE: Using correlation coefficient as the goodness of fit measure, a stopping criterion of Corr > 0.95 is recommended, as it generally reflects a simulated waveform that recapitulates the prominent peaks and troughs of the target ERP. Early stopping of optimization is not currently supported but is under development as this is an important parameter to control. Currently it is recommended to increase the number of epochs if the stopping criteria was not met during an optimization run, but the loss continues to decrease every 10 epochs*.
    4.8.1. If a good fit to the pre-treatment ERP is achieved (i.e. Corr > 0.95), re-adjust the scaling factor to minimize the RMSE and continue to **Step 5**. *NOTE: As described in* ***Step 4.2***, *since “dipole_corr” was applied as the objective function, it is necessary to re-adjust the scaling factor after optimization. In this example, the scaling factor was reduced from the default of 3000x (Figure 7A-C) to 1000x (Figure 7D)*.
    4.8.2. If optimization fails to achieve a good fit to the pre-treatment ERP, return to **Step 4.4** and perform troubleshooting: increase max iterations, find a better manually tuned starting point, identify alternative parameters to adjust. *NOTE: See the section “Troubleshooting when fitting parameters to data features” at the end of the Discussion for a more detailed explanation of troubleshooting steps*.
5. **Establish post-treatment model fit**. Starting with the optimized pre-treatment ERP simulation, manually tune and optimize the **parameters of interest** to fit the post-treatment ERP.
  5.1. Load **post-treatment** empirical ERP waveform from **Step 1** into the graphical user interface (same as **Step 3.1**, Figure 9A).
  5.2. Load the optimized pre-treatment ERP parameters from **Steps 1-4** as a starting point (Figure 9A).
  5.3. Manual hand tuning + parameter optimization (same as **Step 3.1-3.10** and **Step 4**) of **parameters of interest** identified by the user in **Step 1.7** until a high correlation (Corr > 0.95) is achieved. *NOTE: For illustrative purposes, in Figure 9B, hand tuning was used for a signal* targeted *parameter (decreased local network GABA* _*B*_ *maximal conductance), which produced a closer fit to the post-treatment data. However, optimization was not run to see if and how well this parameter change might better account for the data. Rather, the “Representative Results” section below describes how to estimate distributions of multiple parameters hypothesized to be post-treatment* ***parameters of interest*** *using SBI.SBI (detailed in* ***Step 6****) is recommended for rigorous investigations since it estimates the distribution of parameters that account for an ERP waveform, enabling robust comparisons across parameter fits*.
  5.4. **Save model configuration** and compare optimized values for **parameters of interest** between pre-treatment and post-treatment (data not shown).
    5.4.1. Repeat **Step 3.10** to export a .**json** file of model parameters. Move the file to the project folder created in **Step 2.5**.
    5.4.2. View exogenous drive parameters by clicking **Load external drives** (Figure 5A) and select either the pre-treatment or post-treatment network configuration file.
    5.4.3. View local network parameters by clicking **Load local network connectivity** (Figure 9C) and select either the pre-treatment or post-treatment network configuration file.
    5.4.4. Identify changes in parameter values across pre-treatment and post-treatment network configurations. Changes in parameter values correspond to model-based predictions of post-treatment biomarker mechanisms.
6. **Uncertainty quantification with SBI and separability assessment** using the HNN-Python API. *NOTE: SBI requires installation of a separate Python package. See the associated repository https://github.com/ntolley/hnn_jove for a code example detailing how to run parameter inference in HNN with the SBI software package. The code is organized to follow the steps in the subsequent protocol. A full discussion of applying SBI to the HNN model can be found in*^*55*^.
  6.1. Install the SBI package using **pip install sbi**
  6.2. Identify parameter ranges around targeted subset of pre/post-treatment ERP parameters to create a bounded prior distribution for uncertainty quantification
  6.3. Generate training dataset of simulated ERPs with parameters sampled from prior distribution
    6.3.1. Define parameter update function (same as parameter optimization).
    6.3.2. Sample parameters from the prior distribution and generate a dataset of simulations from the parameter samples.
  6.4. Choose a summary statistic that characterizes the EEG waveform (see ^55^ for a complete discussion). *NOTE: A summary statistic is any quantity that characterizes the important features of an EEG waveform. The timing/magnitude of peaks is a common choice. In this manuscript principle component analysis (PCA) is used to extract summary statistics (i.e. the loadings of the first 4 principle components)*.
  6.5. Train the SBI network to match parameter combinations to simulated ERPs *NOTE: The trained SBI network is a python object that can take summary statistics from EEG data as input, and output a distribution of parameters (i.e. the posterior distribution). If successfully trained, simulating these parameters in the HNN model will generate EEG waveforms with similar EEG data (i.e. a posterior predictive check)*.
  6.6. Generate parameter samples and simulations from the SBI network and assess if simulated ERPs sampled from parameter distributions fit the pre/post-treatment ERP
    6.6.1. Provide the experimental EEG waveform as a conditioning input to the trained network.
    6.6.2. Draw parameter samples from the posterior distribution conditioned on the experimental EEG waveform.
    6.6.3. Simulate the parameter samples drawn from the posterior distribution.
    6.6.4. Calculate the similarity between the simulated waveforms to the experimental EEG waveform provided as input. *NOTE: This is known as a posterior predictive check (PPC); a well trained network will produce simulations that are similar to the real waveform (high correlation or low RMSE). If the PPC does not produce simulations that are satisfactorily close to the target waveform, there are two possibilities: 1) the mechanisms considered do not account for the biomarker. In this case new hypotheses must be constructed, and the parameters of the prior distribution must be changed. 2) The SBI network was unsuccessfully trained. In this case, possible solutions include increasing the training budget, and changing the summary statistics of the EEG waveform*.
    6.6.5. If the simulated ERPs from the sampled parameter distributions fit the pre-treatment/post-treatment ERP (PPC with Corr > 0.95), continue to **Step 6.8**. Otherwise continue to **Step 6.7**
  6.7. Troubleshooting SBI network training. *NOTE: A failed PPC indicates the training parameters of the SBI network need to be modified. See the section “Troubleshooting when fitting parameters to data features” at the of the Discussion for a more detailed explanation of troubleshooting steps*.
    6.7.1. Increase the size of the training set.
    6.7.2. Change summary features.
    6.7.3. Choose different SBI architecture for training.
  6.8. Visualize the posterior distribution for the pre- and post-treatment ERPs and assess distribution separability.
    6.8.1. Pass the array of parameter samples from **Step 6.6.2** to the pairplot function and assign the distributions for each ERP a distinct color. *NOTE: The associated code repository demonstrates plotting functionality to reproduce Figure 10*.
    6.8.2. Inspect the diagonal panels of the generated pairplot for non-overlapping distributions and assess the separability of the distributions by calculating the overlap value (OVL, shown in Figure 10A). Parameters with highly separated distributions (OVL < 0.1) correspond to mechanisms of action of the neurotherapeutic that are predicted to change post-treatment relative to pre-treatment. *NOTE: OVL is a metric that quantifies distribution separability in the range of (0,1), with OVL=0.0 indicating no overlap, and OVL=1.0 indicating complete overlap*^*54*, *55*^. *Code to calculate the OVL is provided with the associated code repository*.
7. **Examination/validation and further model constraints**. *NOTE: This step provides examples of how to visualize certain elements of simulated activity in the GUI. These multi-scale details provide targets to validate and inform model derived predictions in follow up experiments*^*7*, *43*^. *This protocol does not provide guidance on how to select which predictions are best suited for validation experiments or how validations experiments should be performed (i.e*. ***Step 7.3***).
  7.1. Load model parameters optimized for pre-treatment and post-treatment and run simulations. *NOTE: The parameters from optimization in Steps 4-5 can be loaded and examined. Examples of how to export network parameters produced by SBI in* ***Step 6****from the Python interface are included in the associated github repository*.
  7.2. Examine multiscale predictions on simulated outputs.
    7.2.1. Plot cell-level spiking activity of optimized simulations (pre-treatment and post-treatment). *NOTE: Certain microcircuit features (e.g. LFP and CSD) are only available through the HNN-Python API. Code based tutorials for all features are documented on the HNN examples page. (https://jonescompneurolab.github.io/hnn-core/stable/index.html)*
      7.2.1.1. Click on the **Visualization tab** (Figure 4A).
      7.2.1.2. Click on the dropdown menu labeled **Layout template** and select **Dipole Layers-Spikes**.
      7.2.1.3. Under the **Dataset** dropdown, select the simulation results to be plotted.
      7.2.1.4. Click **Make figure** to visualize the spiking activity contributing to the dipole waveform.
  7.3. Identify existing datasets and/or collect new empirical data (e.g. invasive electrophysiology, laminar M/EEG, MRS) to test multiscale model predictions.
    7.3.1. If multiscale predictions match empirical datasets, the model is successfully validated for the chosen microcircuit feature.
    7.3.2. If multiscale predictions do not match empirical datasets, update the default HNN network by constraining it to new empirical data and return to **Step 3**.

**Figure 4:**
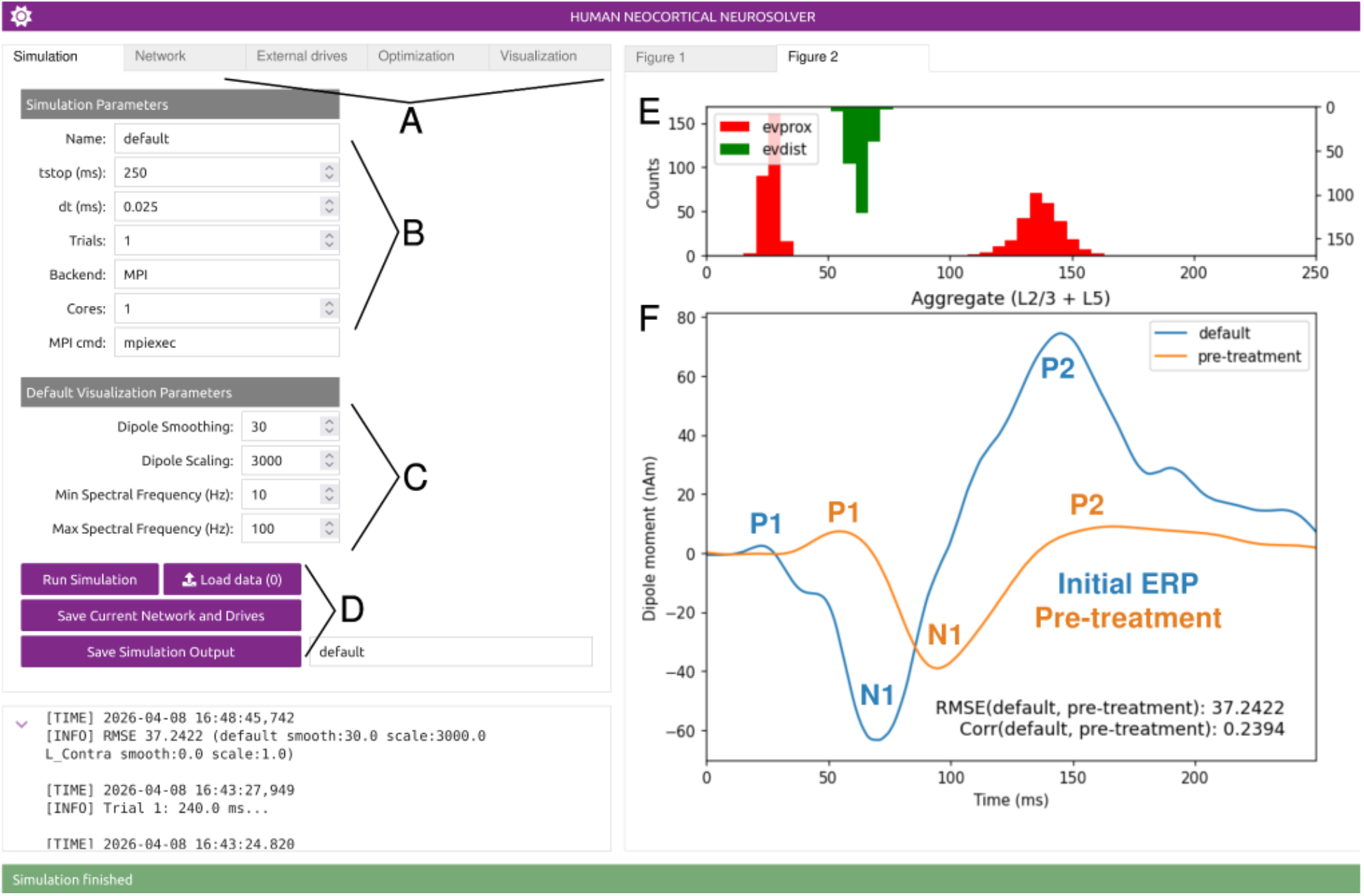
Comparing default canonical HNN simulated ERP waveform to empirical pre-treatment ERP. A: The different categories of model/simulation parameters are accessible through tabs which switch between the different menus. B: The simulation parameters which control settings such as simulation length and number of trials. C: The default visualization parameters which change how simulated waveforms are plotted in the figure panels (E and F). D: The simulation control bar which is used to load data, run simulations, and save model configurations and simulation outputs. E: Spike histograms showing the distribution of exogenous drive inputs for the canonical ERP simulation. F: The dipole waveform of the canonical ERP simulation (“Initial ERP”, blue) overlaid with an empirical auditory ERP (“pre-treatment”, orange) taken from^48^. As shown, the simulation is not a close fit to the data, with peaks that are misaligned in timing and magnitude and Corr <0.95. The following text from Kohl et al., 2022 details the experimental paradigm used to produce the empirical ERP: “Participants were presented with simple 1 kHz sine wave tones, 50 ms in duration with 10 ms fade-in and fade-out time, created in Sound Edit (MacroMedia, San Francisco, CA, USA). The tones were presented alternatingly to the left and right ear (Parviainen et al. 2019). Tones were separated by inter-stimulus-intervals varying between 0.8 and 1.2 s and were presented at 60 dB above the subjective hearing level.”

**Figure 5:**
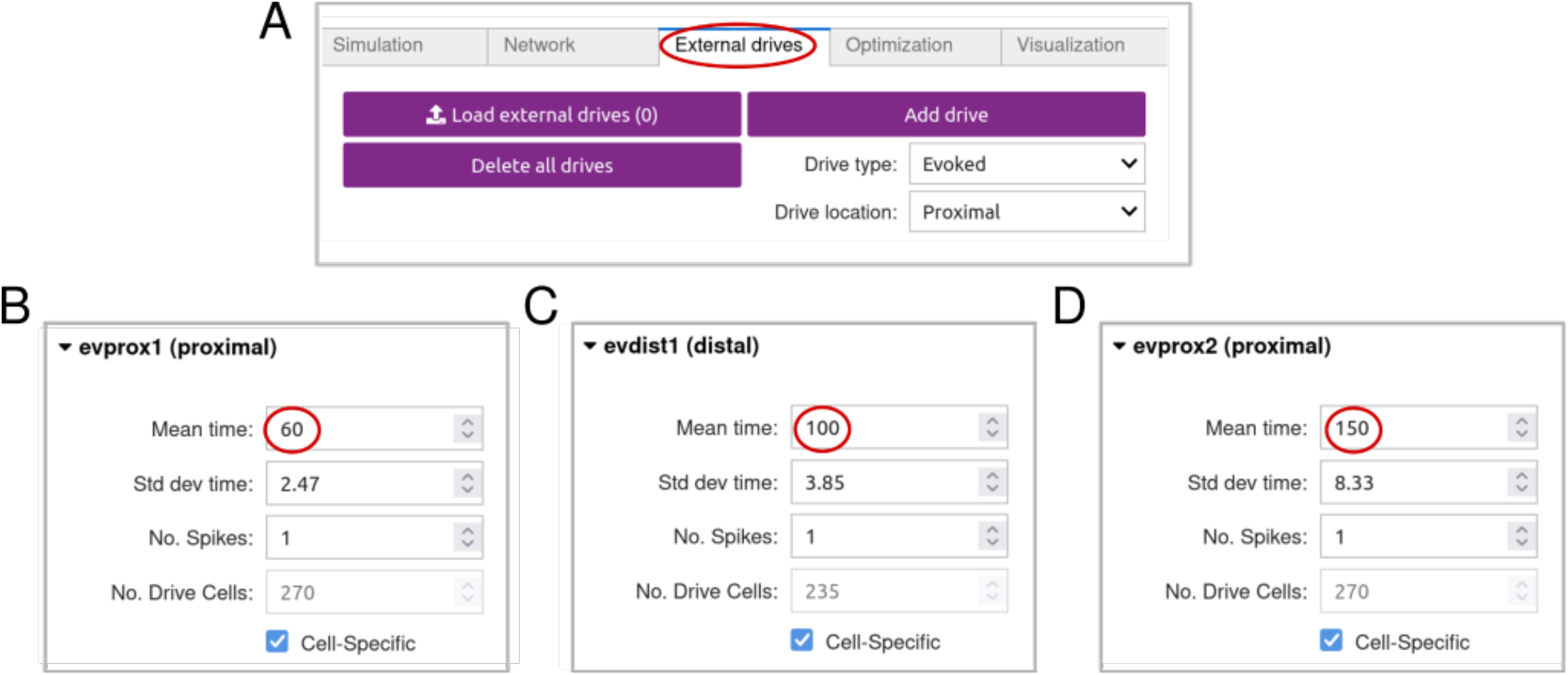
Modifying exogenous drive timing to align peaks. Exogenous drives that activate the local network can be added, modified, and removed through the “External drives” tab. For generating event related potentials, the “Evoked” drive type defines a sequence of proximal and distal drive (see Box 1) whose properties can be specified through parameters that control the spike timing, synaptic location, and synaptic weights. One important parameter for fitting simulated ERPs to data is the **Mean time** (red circle), which can be adjusted to better align the timing of the ERP peaks. Figure 7B shows changing the **Mean time** to align with the ERP peaks at 60, 100, and 150 ms increases the correlation coefficient between the simulation and target waveform.

**Figure 6:**
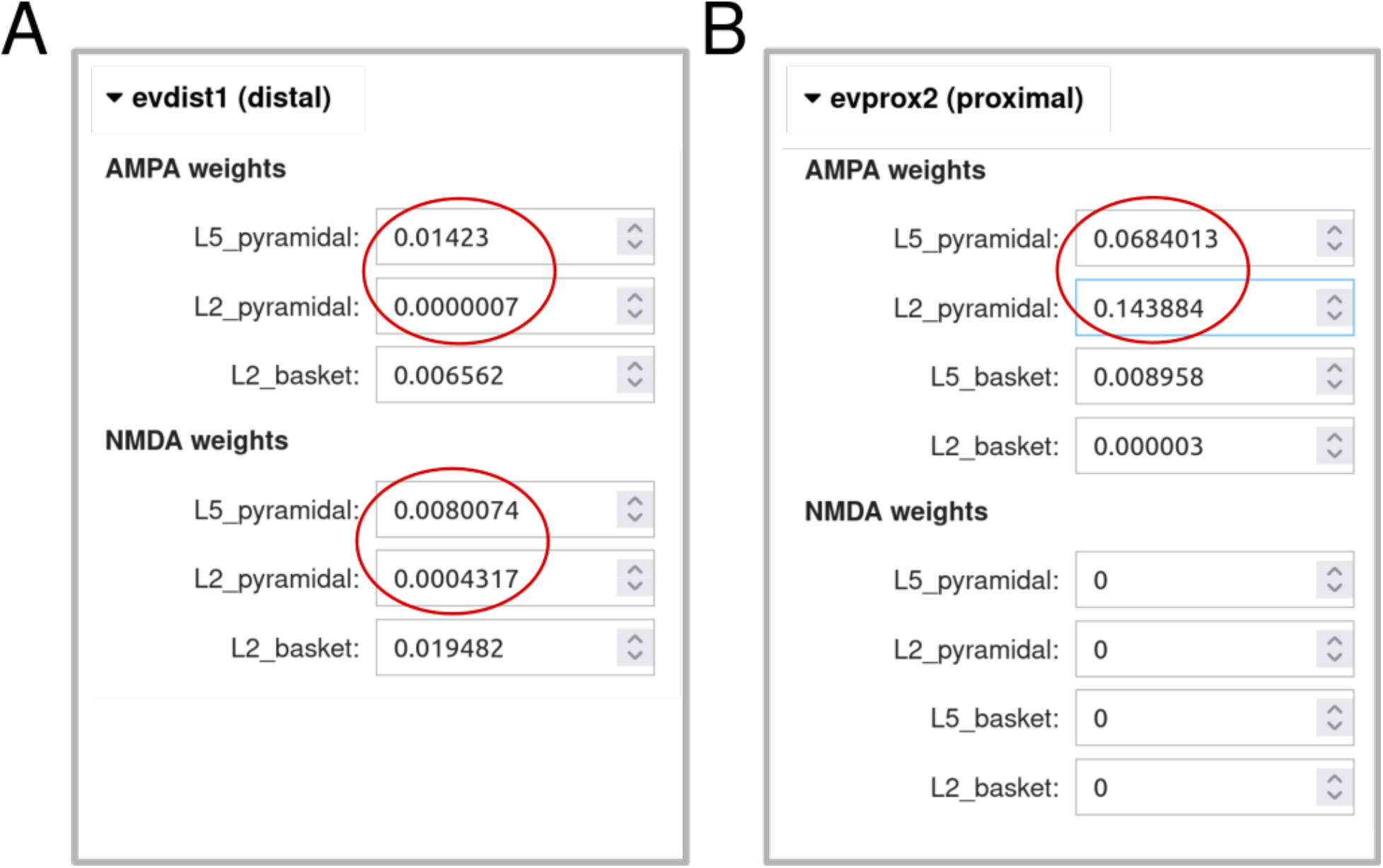
Modifying exogenous drive strength to adjust peak magnitudes. Exogenous drive strengths for AMPA and NMDA synapses can be modified through the “External drives” tab. Synapse weight parameters are located immediately below timing parameters (Figure 5) in the dropdown menu. Boxes for defining the AMPA and NMDA synapse weight onto L2/3 and L5 pyramidal neurons are highlighted in RED circles. In this example, the values were modified to be 10x smaller than the default canonical simulation. Figure 7C shows how this change increases the correlation coefficient between the simulation and target waveform by reducing the magnitude of the N1 and P2 peaks.

**Figure 7:**
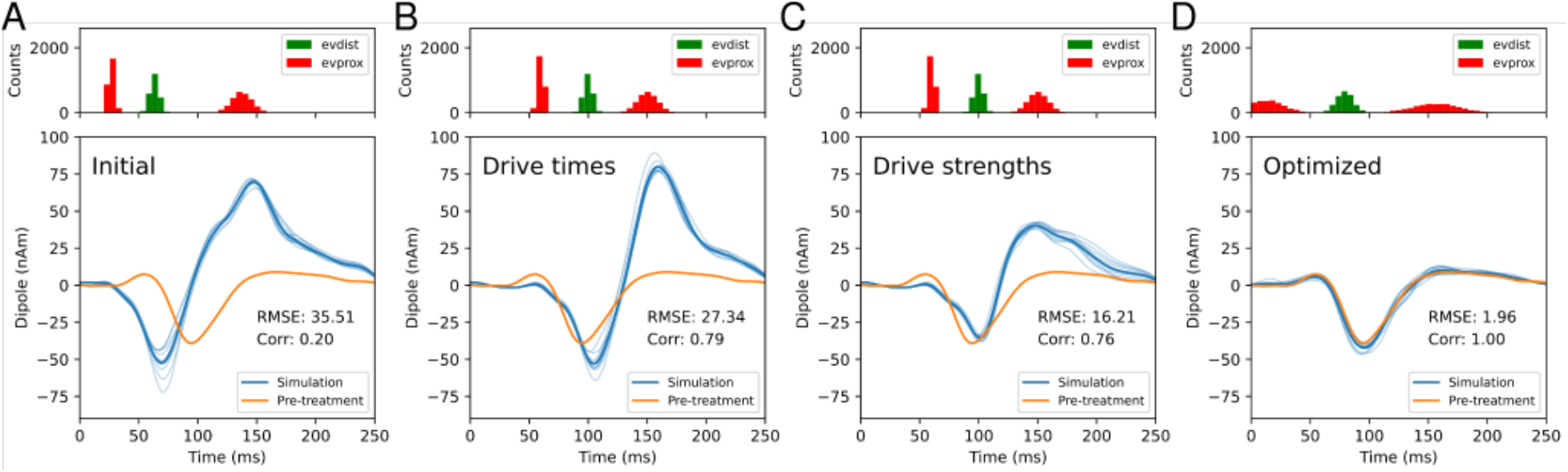
Combining manual hand-tuning and optimization to fit model parameters. The resulting simulations from **Steps 3** and **4**. All simulations show 5 trials with average ERP (dark blue) and individual trials (light blue). A: The initial canonical ERP simulation (blue) overlaid with the pre-treatment auditory ERP (orange). B: Manually adjusting the mean times of the exogenous drives better aligns the peaks of the simulated ERP (blue) to the target waveform (orange). C: Decreasing the AMPA and NMDA strengths of the exogenous input for the first distal and second proximal drive decreases the peak magnitudes of the simulated waveform. D: Automated parameter optimization of the exogenous drive timing and strength produces a simulated waveform (blue) with a very close fit to the target waveform (orange; Corr=1.0). A prominent change is a large increase in the standard deviation of the evoked drives.

**Figure 8:**
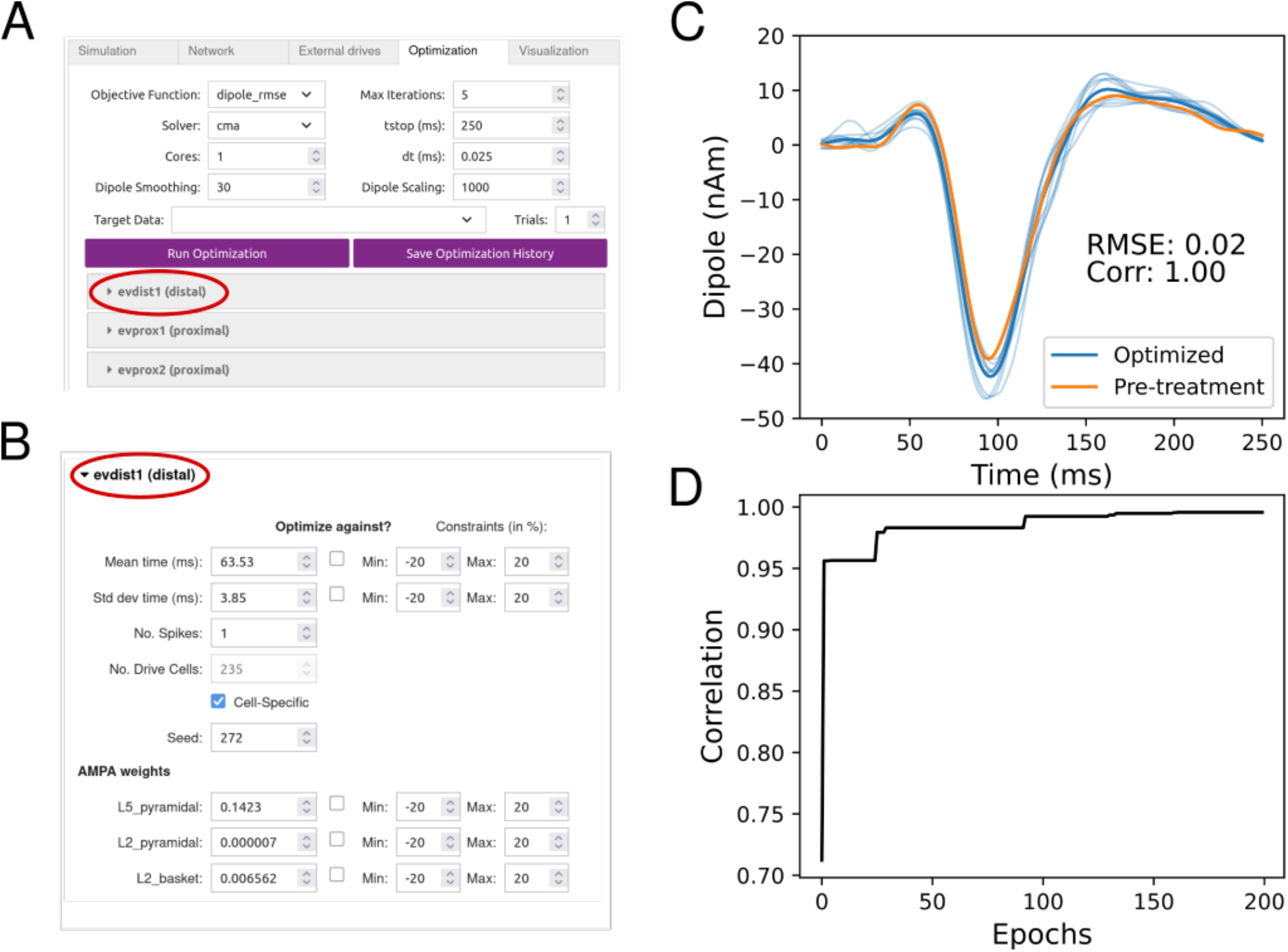
Optimize drive parameters to improve fit to pre-treatment waveform. A: The optimization tab of the GUI provides a simple interface for configuring the settings of a parameter optimization run. B: An optimization run can be customized by selecting which exogenous drive parameters will be optimized, and defining the range of values that can be explored by the optimizer. C: An example of an optimization run where the canonical ERP simulation was fit to an empirical auditory ERP (data from^48^). D: The loss curve for the optimization run in panel C; the best parameter set was identified after approximately 80 epochs.

**Figure 9:**
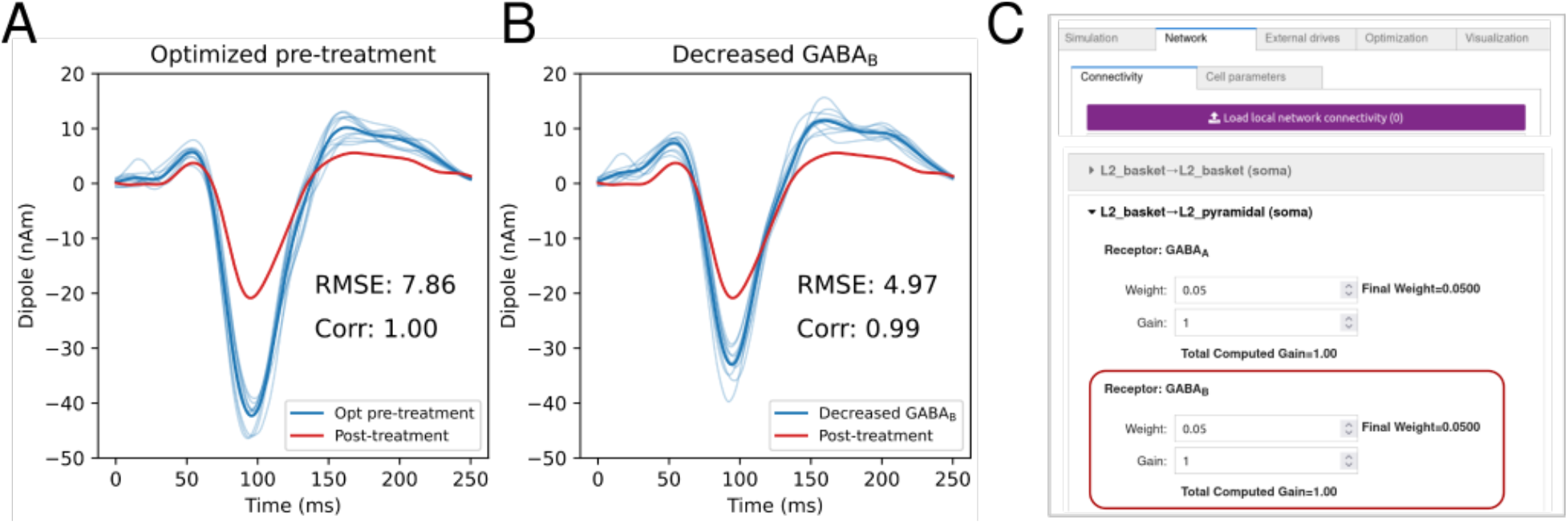
Testing GABA_B_ synaptic strength as mechanism of post-treatment EEG biomarkers. A: The optimized pre-treatment simulations (blue) from Figure 8 overlaid with hypothetical post-treatment EEG (red). The primary difference is reduced magnitude P1, N1, and P2 peaks. B: Decreasing the synaptic strength of all GABA_B_ synapses decreases the N1 magnitude of the simulated ERP, suggesting that the hypothetical neurotherapeutic functions could partially be explained by reducing GABA_B_ signaling. C: Screenshot of the HNN-GUI demonstrating where the synaptic strength of local GABA_B_ synapses can be modified.

**Figure 10:**
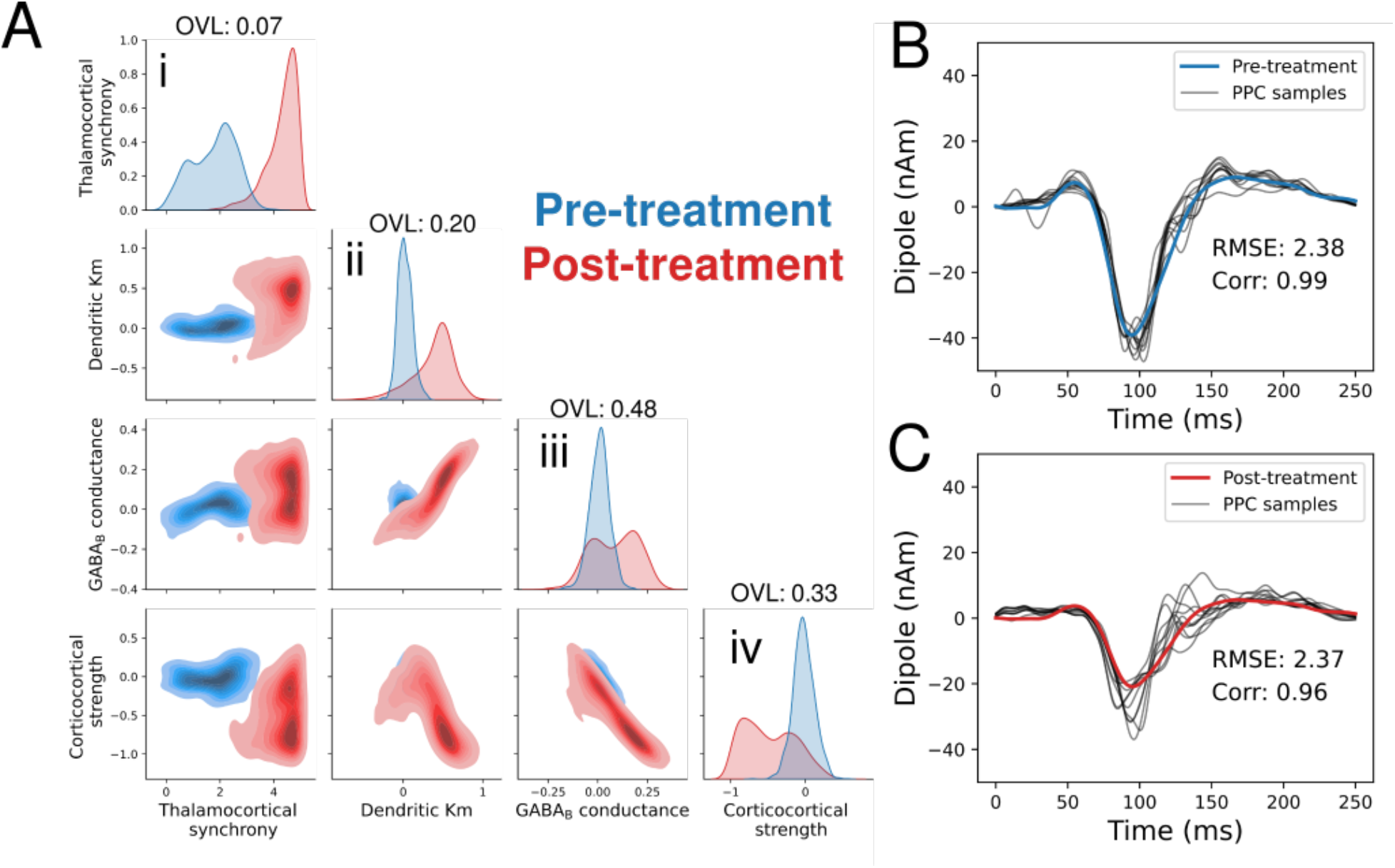
Simulation-based inference for parameter uncertainty quantification to identify distinguishable neurotherapeutic mechanisms generating pre- to post-treatment ERP biomarkers. A: The output of SBI is a multidimensional distribution over parameters, and is visualized using a pairplot. Parameter distributions which account for the pre-treatment (blue) and post-treatment (red) EEG biomarkers show increased values for thalamocortical synchrony (panel i). The low distribution overlap value (OVL=0.07) also confirms that the distributions for thalamocortical synchrony are highly separable. B: The goodness of fit of SBI predicted parameters can be characterized with a posterior predictive check (PPC), where samples from the parameter distribution are simulated and compared to the target waveform. The pre-treatment PPC sampled waveforms (black) closely match the target waveform (blue). C: Same as panel B for the post-treatment ERP. The post-treatment PPC sample waveforms (black) are similarly close to the target waveform.

## REPRESENTATIVE RESULTS

The following section presents a scenario where a neurotherapeutic with an unknown mechanism of action is being investigated with the HNN modeling software. The goal will be to use the pre-treatment and post-treatment EEG signals to make a prediction on how the neurotherapeutic is altering neural circuits. The results shown are entirely for demonstration purposes, with the goal of introducing how HNN modeling is applicable to any neurotherapeutic.

### Developing mechanistic hypotheses underlying EEG ERP biomarker (Step 1)

In this example, a hypothetical sensory event related potential (ERP) paradigm is used to examine how the neurotherapeutic alters the signal (**Step 1.1-1.5**). Figure 1 (top) shows a pre-treatment auditory ERP (blue) alongside a hypothetical post-treatment ERP (red, see also Figure 9). The pre-treatment auditory ERP is experimentally recorded source-localized data from^48^, and the hypothetical post-treatment ERP was generated by scaling the pre-treatment waveform with a gaussian-tapered window. As shown, the hypothetical neurotherapeutic produces a large decrease in the magnitude of the P1, N1, and P2 component relative to the pre-treatment ERP.

*Note that in Kohl et al*., *2022, from which the pre-treatment ERP data was taken, the HNN simulations used a model where the pyramidal neurons were enhanced with more realistic calcium channel dynamics than in the default HNN model. As such, the simulation results in Kohl et al*., *2022 are slightly different from those shown here. The Kohl et al. 2020 model (and other updated and published HNN models) can be accessed through the Python-API (https://jonescompneurolab.github.io/hnn-core/stable/generated/hnn_core.calcium_model.html#hnn_core.calcium_model). Accessing such expanded models though the HNN-GUI is currently under development*.

Next, identify model parameters representing treatment-related effects (i.e.**parameters of interest**) that are hypothesized to explain how the neurotherapeutic reduces the P1, N1, and P2 magnitudes (**Step 1.6-1.8**). Examples of the broad categories of neural mechanisms (and corresponding model parameters) that can be tested include the timing of exogenous synaptic inputs, local neuronal ion channel conductances, local synaptic connectivity, and exogenous synaptic connectivity (Figure 1 middle). This example considers candidate mechanisms from each of these four categories (described below), and uses HNN to test how changes in the corresponding model **parameters of interest** impact the simulated ERP.

#### Parameters of interest

1. **Standard deviation of the first (thalamocortical) proximal drive**(i.e. thalamacortical synchrony),representing variability in the synchronization of initial feedforward sensory inputs.
2. **Muscarinic potassium (Km) channel conductance in L5 pyramidal neurons**, controlling the excitability of the neurons, such that excitability decreases when the conductance is increased.
3. **Local GABA** _**B**_ **receptor strength**, corresponding to a slow time constant inhibitory synapse delivered by interneurons to all cells in the local network.
4. **Conductance strength of subsequent feedback (corticocortical) distal drive**(i.e. corticocortical strength), representing the strength of the ∼100 ms sensory evoked feedback drive via excitatory synaptic input to AMPA and NMDA synapses on cells in the supragranular layers.

### Establishing the pre-treatment ERP model fit (Steps 3-4)

First, simulate the pre-treatment ERP following **Steps 3-4** above (final pre-treatment simulation shown in Figure 8C).

### Establishing post-treatment ERP model fit (Step 5)

Using the pre-treatment ERP as a starting point, hand-tuning and optimization are used to determine if the post-treatment **parameters of interest** can be fit to the empirical post-treatment ERP.

### Uncertainty quantification with SBI (Step 6)

Due to parameter degeneracy that is intrinsic to biophysical models, uncertainty quantification with SBI (**Step 6**) is essential to make predictions on pre-to post-treatment parameter changes. However, a critical step prior to SBI is ensuring that the pre- and post-treatment parameters of interest can be fit to the empirical ERPs (**Steps 3-5**). If this is not possible, then the predictions from SBI will be unreliable, as samples from the posterior distribution will be dissimilar to the empirical waveforms. If a successful fit cannot be found for the pre- and post-treatment ERPs in **Steps 3-5**, then the choice of parameters and their ranges must be changed.

Note that in this example, we only apply SBI to the post-treatment parameters of interest (i.e. the hypothetised treatment mechanisms; 4 in total), and keep all other default and optimized parameters fixed. Applying SBI to a larger set of parameters is possible and provides more robust predictions, however, training with large numbers of parameters is significantly more computationally expensive (see discussion).

To produce robust pre-treatment to post-treatment comparisons, (SBI) can be applied to a set of **parameters of interest** to estimate entire parameter distributions that produce simulations with a close match to the target waveform for both the pre- and post-treatment ERP. In brief, SBI is a Bayesian inference technique which trains a deep neural network to learn the mapping from model outputs to distributions of model parameters (i.e. parameters of interest)^55, 62, 63^. The trained SBI network is then applied to empirical waveforms to identify model parameters that best match the data. A priori hypotheses on the **parameters of interest** and the range of values they can take to produce the waveform are needed. Here, SBI is run over a pre-defined prior distribution of the four **parameters of interest** described above, namely initial thalamocortical proximal drive std deviation (i.e. thalamocortical synchrony), pyramidal neuron dendritic Km conductance, local GABA_B_ conductance, and feedback distal drive conductance (i.e. corticocortical strength). In this example, a uniform prior distribution is used where the minimum and maximum values are calculated as scalar multiples of the default values. Specifically, the prior distribution bounds for thalamocortical synchrony range from 0 to 5 times the default value, and the remaining parameters were 1e-1 to 1e+1 times the default value. Figure 10A shows the results of applying SBI over the hypothesized **parameters of interest** to estimate distributions corresponding to pre- and post-treatment ERPs. Since output of SBI is a multidimensional parameter distribution, Figure 10A visualizes the results as a pairplot. A pairplot organizes projections of the data into univariate distributions on the diagonal panels, and bivariate distributions on the off-diagonal panels. Under this framework, mechanistic predictions correspond to parameters where the distributions between pre-treatment and post-treatment biomarkers are highly separated.

Inspection of the univariate distributions along the diagonal reveals that thalamocortical synchrony has the highest degree of pre-to post-treatment separability (lowest OVL of 0.07), and that it is increased for the post-treatment ERP waveform (Figure 10A(iii) red). In other words, the HNN modeling framework predicts that the hypothetical neurotherapeutic alters the ERP by modifying thalamocortical synchrony (i.e. a distinguishable mechanism of action).

It is important to validate that the parameter distributions produced by SBI actually produce good fitting models of empirical data. One approach is a posterior predictive check (PPC) which draws independent parameter samples from the posterior distribution to run a new set of simulations. The PPC check is considered successful if all randomly sampled simulations accurately fit the target ERP waveform. As shown in Figure 10B-C, the pre-treatment (Figure 10B blue) and post-treatment (Figure 10C red) waveforms closely match the waveforms produced by PPC parameter samples (black), with a correlation coefficient of 0.99 and 0.96 respectively (averaged over 10 independent PPC samples).

### Model Examination and Validation (Step 7)

Using the HNN model, it is possible to directly inspect and visualize the cell- and circuit-level activity such as spiking, underlying each ERP simulation (**Step 7.1**). Figure 11A-B shows a single simulated ERP (black) sampled from the pre-treatment and post-treatment parameter distributions, along with the cell-specific spiking activity contributing the generation of each waveform (Figure 11C-D). The waveform is visualized without smoothing to emphasize the impact of spike timing on the current dipole. In real EEG signals, the electric fields of large populations of pyramidal neurons constructively and destructively interfere producing a signal that appears smoothed at the sensor level. Since HNN models a small cortical region (200 pyramidal neurons), smoothing is used to approximate the effect of simulating larger populations (>100,000) of neurons simultaneously. One notable difference in the pre-treatment and post-treatment spiking activity is observed in L5 pyramidal neurons, such that the amount of L5 pyramidal neuron spiking decreases post-treatment relative to pre-treatment (Figure 11C-D red dot; note that Figure 11 only shows spiking for a single sample from the posterior distribution, to produce testable predictions, multiple samples should be drawn and simulated). All together, it can be seen that the hypothetical treatment-related effects alter the multi-scale circuit activity to decrease the magnitude of the P1-N1-P2 peaks.

**Figure 11:**
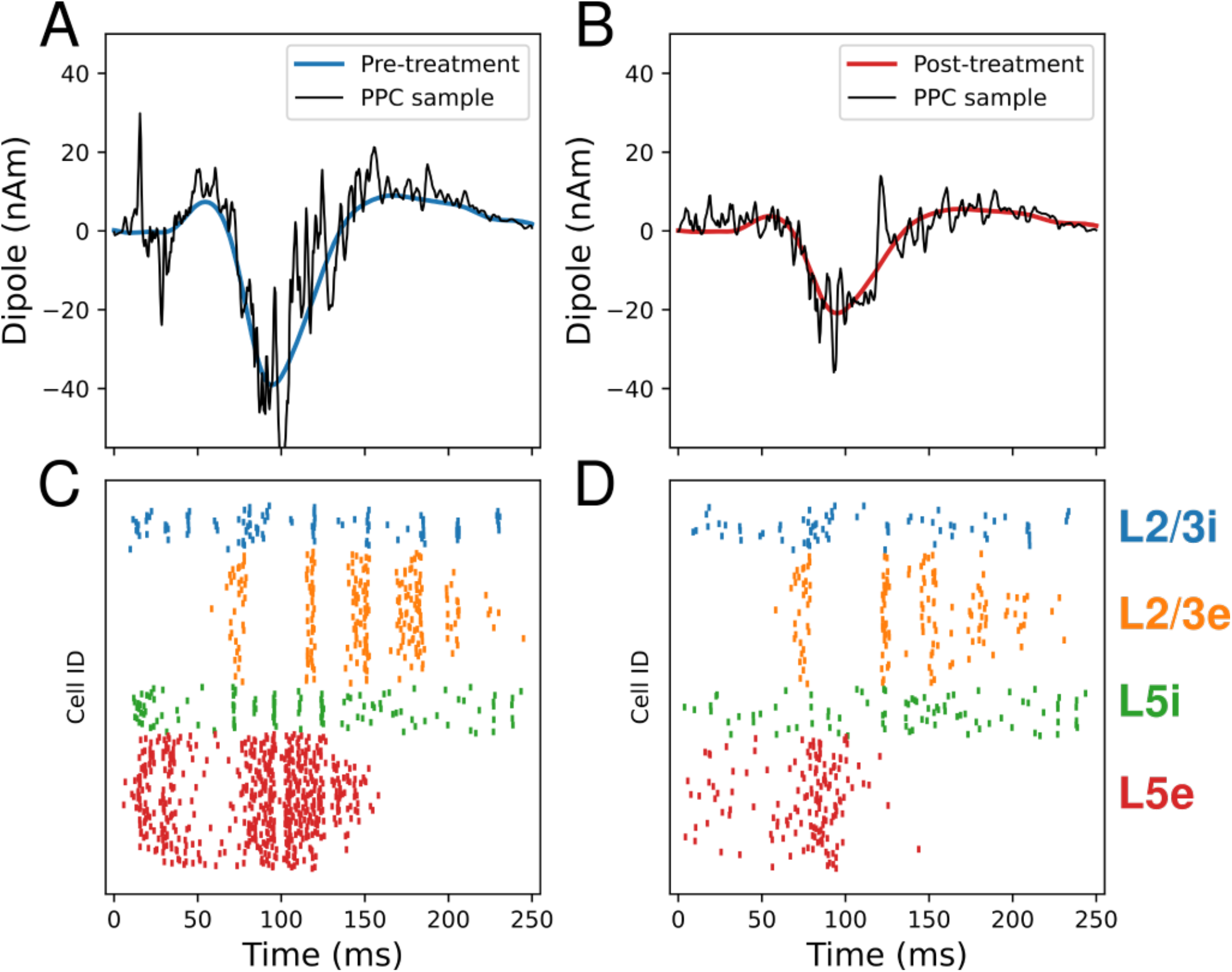
Cell-level spiking activity contributing to generation of EEG biomarkers. A: The ERP waveform for the pre-treatment condition (blue) with a single simulation from model parameters fit using SBI (black). The simulation represents a single posterior predictive check (PPC) sample from the pre-treatment parameter distribution in Figure 10A (blue distribution), and is plotted with no smoothing. B: Same as panel A for the post-treatment ERP waveform (Figure 10A orange distribution). C: The simulated spiking activity underlying the pre-treatment PPC sampled ERP waveform. D: Same as panel C for the post-treatment ERP waveform.

Predictions such as these can be directly tested through invasive electrophysiology (e.g. with high-density laminar probe electrodes^64^) in animal models. As multi-scale information is obtained from invasive recordings (or through other imaging modalities), these data can be used to further constrain the model-derived predictions. Although this protocol shows how to fit the HNN model to macroscale EEG and then infer underlying microcircuit activity, a workflow can also be developed to perform the reverse: fitting to microcircuit activity (cell specific spiking, LFP/CSD) and then inferring the corresponding macroscale EEG.

## DISCUSSION

Computational neural modeling of EEG biomarkers may allow deeper insight into how CNS therapeutics reconfigure neural circuits and provide predictions on what biological processes may be related to therapeutic effects. The workflow above demonstrates how a commonly measured EEG biomarker, auditory event related potentials, together with biophysical modeling with HNN, can be used as a window into the mechanisms by which a drug impacts neural activity. The overall strategy can be adapted to investigate other local EEG signals, including low-frequency neural oscillations^45, 65^, and transient spectral events^7, 43, 66^, see Box 1.

Compared to other frameworks for neural modeling of EEG, HNN (detailed below) offers a balance of model complexity and computational efficiency. For example, The Virtual Brain (TVB) is an EEG modeling framework that can account for brain-wide networks that generate spatiotemporal EEG signals^34, 67^. To do so, the models of neural activity are mathematically reduced, eliminating cell details (e.g. pyramidal neuron morphology) and limiting the ability to predict drug effects at the cellular level. Alternatively, large-scale morphologically and physiologically detailed models are able to simulate EEG signals with more realistically detailed neurons compared to HNN^68–71^, however this detail comes at an extreme computational cost (i.e. several hours of compute time to simulate a few seconds) that makes interpretability and hypothesis testing challenging for general scientists. HNN’s intermediate complexity (Figure 2) permits the ability to make localized cell- and circuit-level predictions with moderate computational resources (i.e. seconds to simulate a few seconds).

There are several limitations to using EEG and biophysical neural modeling to study brain disease and drug mechanisms. Importantly, the biophysical cell and circuit properties generating EEG signals may not fully encapsulate the array of drug and disease effects, with many important properties such as immunologic responses potentially not directly impacting EEG, and hence undetected in such experiments. Further, the scientific basis for biological mechanisms is often motivated by animal models, which are imperfect representations of human brain circuits and may map poorly onto, for instance, neuropsychiatric conditions where symptoms are based on neuropsychological assessment^72, 73^. Another challenge is untangling the mechanisms of acute vs. chronic pharmacological effects. While the biochemistry and physics dictating drug-receptor interactions provide highly accurate models of acute drug effects, less is known about the sustained biological changes induced by the same drugs. A specific limitation of HNN is that the model represents a single localized canonical neocortical network, while neurotherapeutics and CNS disease likely impact regions across the entire brain. Effects of drugs on downstream circuits that drive the HNN model can be represented by changes to the timing and strength of exogenous drives. However, direct empirical examination of how drugs/diseases modify cells and local connectivity in the exogenous regions is not always feasible.

Parameter degeneracy is another fundamental challenge in all biophysical models of neural systems, as multiple parameter sets can produce nearly identical solutions. We show that SBI is useful for this challenge as it characterizes distributions of parameters that produce similar solutions (Figure 10). While SBI enables quantification of parameter uncertainty, for computational tractability, it was applied in this protocol to a limited set of parameters corresponding to drug mechanism hypotheses. The assumption made on the non-estimated parameters influences the resulting dynamics of the network. One approach to expand the estimation process to a larger set of parameters is a form of SBI known as sequential neural posterior estimation (SNPE), which sequentially refines parameter estimates for a single waveform, facilitating estimation of high-dimensional parameter distributions (>10 dimensions). Beyond quantifying parameter degeneracy with SBI, directly constraining the model to additional experimental data can help limit the possible parameter space and permit more precise model-based predictions. Because the primary generators of EEG signals reflect dynamics across the cortical layers, tools such as invasive laminar electrophysiology with microelectrodes (i.e. cell spiking, LFP, and CSD) can be powerful pieces of information for narrowing the range of plausible EEG mechanisms.

As EEG technologies continue to be integrated into preclinical research, clinical trials, and clinical decision-making settings, mechanistic understanding will be essential for linking therapeutic mechanisms with the biological basis of CNS diseases across cohorts and within individuals. Results from computational modeling and simulation (CM&S) are being increasingly recognized by federal regulatory agencies as viable evidence for the safety and efficacy of medical interventions^74, 75^. Further, CM&S studies advance the principle of the 3R’s (reduce, refine, replace) for ethical animal research, as they can be used to design targeted validation experiments (reduce and refine) and derive maximal value from animal use with sufficiently mature computational models^76, 77^. Realizing the full potential of biophysical simulation tools like HNN will require sustained validation against real data across physical scales, experimental paradigms, and contexts of use, such as validating model predictions with ground truth signals from invasive recordings. Such validation, as well as the uncertainty quantification method covered by this protocol, may serve as critical components of future efforts to meet verification, validation, and uncertainty quantification (VVUQ) standards adopted by the FDA and other regulatory and industry stakeholders to assess the credibility of neural circuit models ^74^.

### Critical steps for protocol success

Once you have an established ERP biomarker and have properly installed the software (**Steps 1-2**), the critical steps to successfully going through the protocol are getting to a “Yes” in each of the boxes in the flowchart (Figure 3). If you end up at a “No”, there are alternative procedures to follow. Success in Steps 3-5 requires an initial choice of hypothesized parameters that can be optimized to achieve a good fit between simulated and recorded pre-treatment/post-treatment ERPs. If you end up with a “No”, the user should identify alternate parameters and test that they can be optimized to fit the data at hand. The likelihood of getting repeated failures (“No’s”) is low given prior success fitting the model to data, but if the user does get repeated No’s then the default HNN network model parameters will need to be modified, and/or further biophysical detail will need to be added. Success in **Step 6** requires a proper choice of several SBI features, and if you end up with a “No” then there are explicit troubleshooting steps discussed below. At the end of **Step 6**, you will have successfully followed all protocol steps outlined in the manuscript, arriving at testable model-based predictions and uncertainty quantifications.

**Step 7** is critical to validation of model-based predictions, however the exact procedures are beyond the scope of this manuscript and depend critically on the empirical modalities that the user has access to. Some example possible validation procedures include, laminar electrophysiological recordings that can test prediction on layer and cell specific spiking patterns and/or LFP and CSD (e.g. see^7^), layer resolved E/MEG (e.g. see Bonaituo et al 2021), magnetic resonance imaging (MRI) spectroscopy and/or positron emission tomography (PET) scans that can provide a measure of GABA or glutamatergic tone, and diffusion tensor imaging (DTI) that can provide insight into thalamocortical fiber integrity.

### Troubleshooting and customization

#### Troubleshooting when fitting parameters to data features

**Steps 3-6** of the protocol concern techniques to fit model parameters to data, here we provide potential issues encountered during these steps and proposed solutions.

##### Parameter optimization (**Steps 4-5**) is not converging to a high correlation coefficient

If parameter optimization does reach a high correlation coefficient Corr > 0.95, several modifications can be made including 1) adjust optimizer hyperparameters, 2) adjust scaling factor and smoothing parameters, and 3) adjust exogenous drive parameter ranges to consider, and/or add additional external drives. For optimizer hyperparameters, using the CMA-ES solver with a large population size (>100 simulations per epoch) will increase the robustness of the search, however, the computational cost will increase, leading to potentially long wait times. For the scaling and smoothing parameters, a poorly chosen smoothing value of the simulated waveform may severely hinder performance, and can be addressed with multiple optimization runs over a range of smoothing parameters (between 5-60 ms). For exogenous drive parameters, expanding the min/max bounds of each parameter may be necessary if the initial simulated ERP is far from the experimental waveform. It may also be necessary to increase the number of proximal/distal drives, as the proximal-distal-proximal pattern of ERPs was established for evoked responses in primary sensory cortices, and may change depending on the stimulation paradigm and brain region being modeled.

##### Optimized simulations **(Steps 4-5)** present a high correlation but are missing ERP features present in the experimental data

Since correlation based loss functions show the largest decrease for high magnitude peaks in ERP, small magnitude peaks like the P1 may be missed entirely despite an apparently high correlation coefficient. Two modifications to address this are 1) set a stricter loss threshold, or 2) multiply the time periods with small magnitude peaks with a weighting factor; this is applied to both the simulated and experimental waveforms so that these features have a larger contribution to the loss.

##### SBI parameter distributions produce simulations that are dissimilar to the target waveform (failed posterior predictive check **Step 6.6**)

The first check will be to plot the simulated waveforms used to train the SBI model (i.e. Figure 10B-C, code to produce plot included in code repository). If all simulations are highly dissimilar from the experimental ERP (large RMSE or correlation Corr < 0.95), then prior distribution needs to be modified to either include new parameters, or expand the bounds of existing parameters. Increasing the size of the training set may also be necessary depending on the number of parameters optimized. For amortized inference (i.e. an SBI network that is trained once and can be re-used for multiple experimental waveforms), a rule of thumb is 10^N^ simulations for N parameters. For non-amortized inference (i.e. an SBI network that is trained to produce the posterior distribution for a single experimental ERP), a significantly larger number of parameters can be optimized. Alternative approaches to improving training of the SBI network include changing the summary statistic features and changing the SBI network architecture.

#### Modifying default network to include alternative or additional circuit details

The field of biophysical neural modeling of EEG signals is still in its infancy, and how much detail is needed to make accurate predictions on cell and circuit elements contributing to localized ERP generation is still largely unknown. As described in the “Iterative process for using HNN section”, HNN’s framework is built around established biophysical principles of canonical neocortical circuitry and has proven to make generative and testable predictions on multiscale elements that contribute to EEG in several sensory and frontal brain areas. However, users may want to investigate the impact of other cell or circuit elements not included in the default model on simulated ERPs, for example additional interneuron types or further cortical layer details. Adding additional circuit details constrained by data becomes essential when the validation step in the protocol (**Step 7**) fails. HNN is designed in a modular fashion to accommodate such modification. Properties such as cell biophysics and synaptic connectivity can be modified within HNN’s graphical user interface, while modifications to more advanced properties such as cell morphology, number, and location require use of the Python interface. As one example, in Diesburg et al., 2024^51^ the default HNN model was modified through the Python interface to account for more specific interneuron connectivity structure in frontal cortices. Updated model simulations lead to further testable predictions for testing and validation in follow up empirical studies. HNN’s open framework facilitates the sharing of expanded models for others to use and build from.

## ACKNOWLEDGMENTS

All code used to produce the results shown in this protocol can be found at https://github.com/ntolley/hnn_jove

This work was supported by the Brown Biomedical Innovation to Impact Award, the National Institute of Health (NIH; https://www.nih.gov; grant numbers U24NS129945 and P20GM130452), and the National Science Foundation (NSF; https://www.nsf.gov; grant number 2424101). The funders had no role in study design, data collection and analysis, decision to publish, or preparation of the manuscript.

